# Glycolysis and hexosamine biosynthesis pathways are key for inflammatory protein maturation and leukocyte adhesion to human aortic valve cells

**DOI:** 10.64898/2025.12.12.693904

**Authors:** Tania Sánchez-Bayuela, Mirian Peral-Rodrigo, Javier López, Cristina Gómez, Enrique Perez-Riesgo, Natalia López-Andrés, Nieves Fernández, J. Alberto San Román, Mariano Sánchez Crespo, Carmen García-Rodríguez

## Abstract

Inflammation and metabolism reprogramming are hallmarks of calcific aortic valve disease (CAVD). Recent studies link inflammation to hyperglycolysis and calcification in valve interstitial cells (VICs). The metabolism of valve endothelial cells (VECs) has received less attention despite both resident valve cells are exposed to alike inflammatory clues involved in the biosynthesis of pathologically relevant glycoproteins during the early stages of CAVD. On this basis, we investigated the outcomes of glucose metabolism rewiring on glycoprotein maturation and immune cell adhesion in human resident valve cells. Real-time metabolic analysis revealed that basal VECs are more glycolytic than VICs. Also, VECs and VICs exposed to inflammatory stimuli exhibited a distinct rewiring, with VECs shifting to a more energetic metabolism, despite a similar upregulation of glycolytic genes. Blunting glucose metabolism in VICs and VECs inhibited inflammatory routes canonically associated with glycolysis, and the expression of proteins associated to the inflammatory response like interleukin-6 and cyclooxigenase-2. Moreover, Western blot and adhesion assays revealed that glycolysis is necessary for the expression and post-translational modifications of intercellular adhesion molecule-1 and vascular cell adhesion molecule-1, and the ensuing process of monocyte-VECs adhesion. Notably, inhibition of the hexosamine biosynthetic pathway using DON and of N-glycosylation by tunicamycin, further disrupted adhesion molecule maturation and monocyte-VECs adhesion. In conclusion, glycolysis and its side-branch route the hexosamine biosynthesis pathway are necessary for nutrient-driven post-translational modifications of inflammatory proteins in inflamed valve cells and the subsequent process of monocyte-VECs adhesion that plays a key role in the initial stages of CAVD pathogenesis.

**NEW & NOTEWORTHY:** The study uncovers a relevant role of glycolysis and its side-branch route the hexosamine biosynthesis pathway in sugar-driven post-translational modifications that are critical for the proper function of leukocyte adhesion molecules and other relevant proinflammatory molecules in aortic valve cells. These events are essential for the recruitment of cells of the monocytic lineage to aortic valve leaflets in the initial stages of calcific aortic valve disease.

**GRAPHICAL ABSTRACT:** 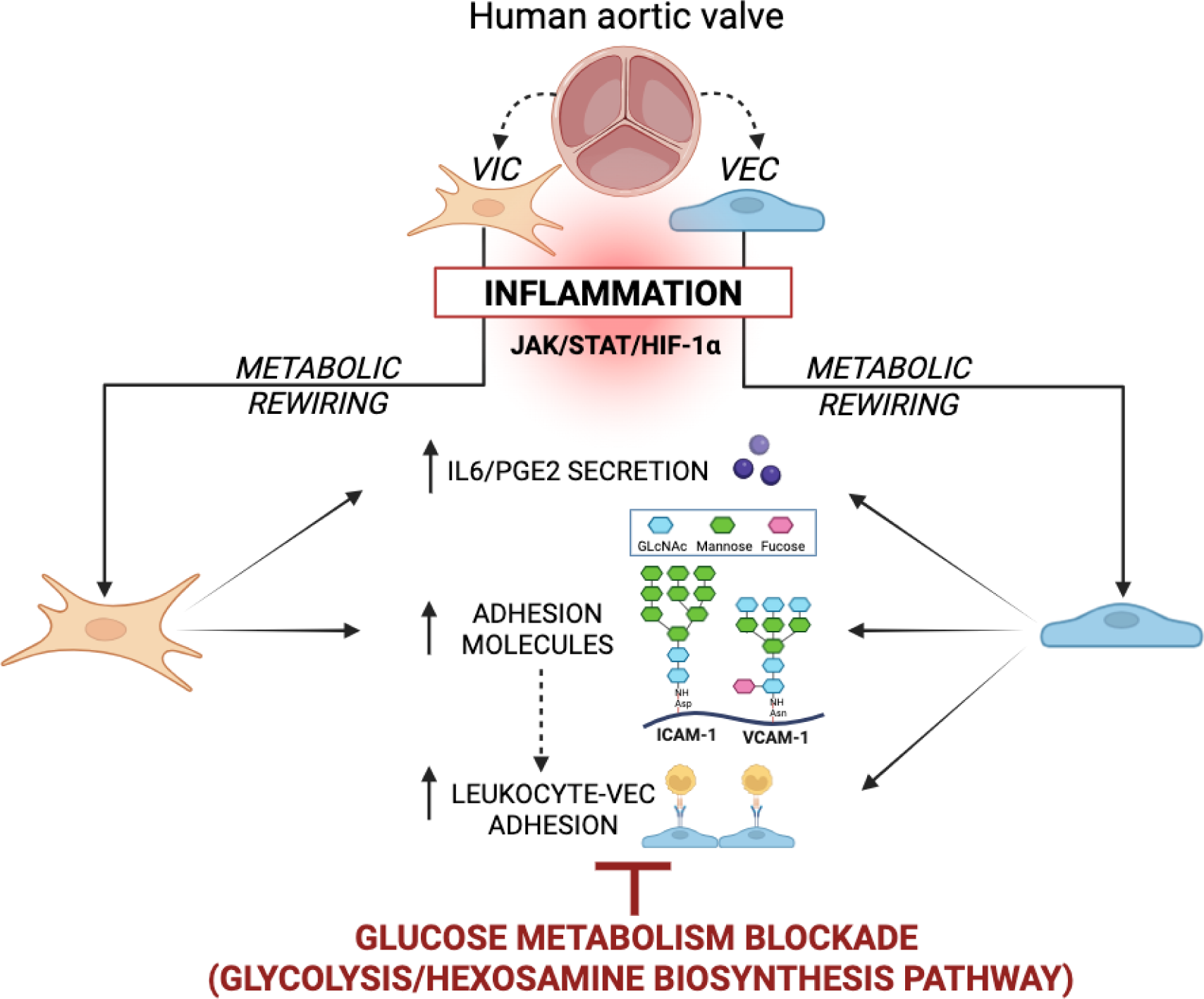

## INTRODUCTION

Inflammation and metabolic reprogramming are hallmarks of calcific aortic valve disease (CAVD), the most prevalent valvulopathy characterized by a progressive fibrocalcific remodeling that impairs valve function and heart function (1, 2). Accumulating evidence supports inflammation as a driving force in CAVD pathogenesis (3–5). Inflammation mediated by the axis Janus kinase (JAK)-signal transducer and activator of transcription (STAT)/hypoxia inducible factor (HIF)-1α promotes the production of cytokines and adhesion molecules, which entail adverse and sex-differential consequences on resident valve cells (6–8). Among the effector molecules are the cytokines interleukin (IL)-6 and IL-8, and the enzyme cyclooxygenase-(COX)-2, involved in the synthesis of prostaglandin E_2_ (PGE_2_) (3, 9). Cytokines, COX-2, and adhesion molecules are glycoproteins playing pathophysiologically relevant roles in CAVD (9–11), including the processes of calcification and leukocyte adhesion to valve endothelium.

Glucose metabolism is influenced by the inflammatory response in vascular and cardiac cells (12, 13). While healthy human cardiomyocyte primarily rely on fatty acid oxidation, a shift towards glycolysis is associated with heart failure (14). In this connection, the metabolic rewiring on CAVD is gaining increasing attention (15–18) and recent work links inflammation, hyper-glycolysis, and calcification in aortic valve interstitial cells (VICs) (17). Moreover, primary cultures of VICs exposed to inflammatory conditions mimic the metabolic alterations observed in VICs from CAVD patients (17). A key driver of this rewiring is HIF-1α, the stabilization of which by hypoxia and inflammatory signals regulates gene transcription (19).

Glycosylations are sugar-dependent post-translational modifications (PTM) critical for protein functions. They are essential for cell adhesion and glycosylation deficits may disturb proper inflammatory responses (20, 21). The hexosamine biosynthetic pathway (HBP), an offshoot of glycolysis, yields the activated amino-sugar uridine diphosphate N-acetylglucosamine (UDP-GlcNAc), which together with other nucleotide sugars, are the building blocks for glycoprotein biosynthesis (22). Glutamine fructose-6-phosphate amidotransferase (GFAT) is the rate-limiting enzyme of HBP that controls the entry of glucose-derived carbons into this pathway (23). Altered glycosylation patterns can entail pathological processes, and the HBP integrates glucose, glutamine, acetyl-CoA, and other metabolites to yield UDP-GlcNAc to drive glycosylation and protein maturation.

In this study, we addressed the role of the metabolic rewiring of human VECs and VICs in the maturation of glycoproteins and the ensuing consequences of this rewiring in the adhesion of immune cells and the production of proinflammatory molecules involved in the initial stages of CAVD.

## METHODS

A detailed version of the methods including the Major Resources Tables and the study design is available in the Supplemental Material.

### Human aortic valve tissue and valve cell cultures

The aortic valve leaflets were collected at the Hospital Clínico Universitario de Valladolid after obtaining written consent from patients. The study was conducted in accordance with the Helsinki Declaration and was approved by the Institutional Review Board (IRB protocol numbers PI 15-263 and PI 21-2403). Aortic valves were obtained from excision of tricuspid aortic valves leaflets from N=36 heart transplant recipients with no valve disease (31 males/5 females, 60.19 ± 5.32 / 59.8 ± 7.01 years, *p*-value= 0.88, Supplemental Table S1). European guidelines for indications for heart transplantation were followed. Inclusion criteria for non-stenotic aortic valves were heart transplant recipients with no valve disease, which was excluded by echocardiographic evaluation. Resected leaflets were used to generate VICs and VECs cultures using sequential digestions with collagenase II as previously described (8, 24).

*Ex vivo* studies were performed on surgical aortic valve samples from CAVD patients undergoing surgical valve replacement at Hospital Universitario de Navarra (Pamplona, Spain). The study was approved by the Research Ethics Committee (IRB protocol 2013/26, n. 137) and conducted in compliance with the Declaration of Helsinki. All recruited patients provided informed written consent. Clinical and demographic data of the aortic valve donors included in this study have been previously reported (25).

### Cell activation and pharmacological treatment

Cells were activated in M199 medium-2% FBS (activation medium) for 24 h. Stimuli included recombinant interferon (IFN)-γ and bacterial lipopolysaccharide (LPS) from *Escherichia coli* O111B4. In pharmacological studies, 2-DG, D-(+)-mannose, and tunicamycin were incubated for at least 30–45 min prior to activation.

### Real-time metabolic analysis of human VECs and VICs

The metabolism of VECs was examined in an Agilent Seahorse XFe24 metabolic analyzer (Agilent Technologies, Santa Clara, CA) using Cell Mito Stress and ATP assay kits. In total, 40 x 10^3^ VECs or 30 x 10^3^ VICs were seeded in a Seahorse XF24 Cell Culture Microplate and allowed to attach overnight. Then, cells were activated as indicated for 24 h. Before assay, media were replaced by Seahorse XF RPMI Medium, pH 7.4, supplemented with 2 mmol/L L-glutamine, 25 mmol/L D-glucose, and 1 mmol/L pyruvate. Cells were incubated for at least 45 minutes at 37°C in a non-CO_2_ incubator before the analysis. Data were analyzed with Seahorse Analytics software and normalized to the number of cells, evaluated by Hoechst staining in a Cytation 5 multimode reader (BioTek, Winooski, VT) at the end of the experiment. Data describe the indicated biological replicates performed in triplicate.

### Reversed transcriptase quantitative PCR (RT-PCR)

Total RNA was extracted with Tri-Reagent and analyzed by RT–PCR as previously described (6, 17, 26). For the experiments, one well of a 6-well plate of cultured cells were used. Specific primer sequences are listed in Supplemental Table S2. First-strand cDNA, synthesized by the reverse transcription reaction, was later amplified by PCR in a LightCycler480 (Roche Diagnostics, Rotkreuz, Switzerland) using KAPA SYBR FAST master mix kit (Merck Millipore, Burlington, MA). Transcript levels were expressed as relative to the housekeeping gene value, *GAPDH* (2^−ΔCt^, Ct=cycle threshold value). Data describe the number of biological replicates performed in duplicate.

For aortic valve tissue from CAVD patients, total RNA was isolated according to a standardized phenol-chloroform protocol, using Qiazol reagent and miRNeasy mini Kit (QIAGEN), and reverse-transcribed into single-stranded cDNA, using an iScript Advanced cDNA Synthesis Kit (Bio-Rad). Downstream qPCR amplification of first-strand cDNA was performed using iQ SYBR Green Supermix (Bio-Rad) in a CFX Connect Real-Time PCR System (Bio-Rad). The relative expression of each selected gene product was calculated using the 2^−ΔΔCt^ method by normalizing the target’s readout to the Ct readout of the housekeeping genes *ACTB*, *18S*, *GADPH*, and *HPRT*.

### Immunodetection of adhesion molecules and signaling routes by Western blot

Whole cell lysates were analyzed by Western blot as previously described (6, 24). Densitometry data were presented as arbitrary units normalized to β-tubulin. Primary antibodies used in the study included p-STAT1, p-NF-κB, HIF-1α, intercellular adhesion molecule (ICAM)-1, vascular cell adhesion molecule (VCAM)-1, GFAT2, and β-tubulin antibodies (Major resource table). Uncropped blots are shown in the Supplemental information. Data refer to the indicated biological replicates.

### Quantitation of interleukin IL-6 and PGE_2_ secretion by ELISA

Cytokine secretion was detected in the supernatants of cells activated for 24 h by immunoassay kits following the manufactureŕs procedures: IL-6 (Diaclone, Besançon, France, # 950.030.048) and PGE_2_ (Arbor assay Inc, Arbor Assay, MI, #K051-H1). The absorbance at 450 nm was measured using a microplate reader (Versamax, Molecular Devices), and data were normalized to total protein content measured by the bicinchoninic acid/BCA method.

### Monocyte-VECs adhesion assays

VECs monolayers cultured in a 6-well plate were pre-incubated with metabolic modulators and then activated in activation media for 24 h under static conditions. In parallel, a monocytic cell line, THP-1 (ATCC®; Middlesex, UK Ref. TIB-202TM), was cultured in RPMI-1640 medium supplemented with 1% L-glutamine and 10% inactivated fetal bovine serum. For the adhesion assay, 1 x 10^6^ cells/well of THP-1 in HBSS containing Ca^2+^ and Mg^2+^, were added to VECs monolayers and incubated for 7 min at 37°C. After two washes with HBSS, microphotographs of 16 sections were taken using a phase-contrast microscope (Nikon Eclipse TS100) connected to a digital camera. The number of adhered THP-1 cells was quantified using Image J software. Data were expressed as total number of THP-1cells adhered to the area of each section (mm^2^).

### Immunohistological Evaluation

Tissue staining was performed on transversal sections of human aortic valve leaflets and analyzed as described (25). All grossly calcified valves were decalcified in 10% formic acid solution (Sigma) for 24 h. Samples were dehydrated, embedded in paraffin, and cut in 5-μm-thick sections.

Immunohistochemistry was performed following the protocol of Leica BOND-Polymer Refine Detection automatic immunostainer (Leica). All solutions were filled into the bottle-Bond Open Container (Leica) and registered on computer using the Leica Biosystem program. The immunostaining program protocol includes Fixative solution, Bond wash solution, Blocking with common immunohistochemistry blocker, and incubation with the primary antibody for ICAM-1 or VCAM-1 (Santa Cruz Biotechnology). Secondary antibodies poly-HRP-anti-mouse or poly-HRP-anti-rabbit IgG were used as appropriate. Positive immunoreactive signal was developed using an enhanced 3,3’-diaminobenzidine (DAB) system (Leica). Slides were counterstained with hematoxylin and mounted using DPX mounting media (Merck/Sigma-Aldrich). Imaging was performed using a Leica microscope at 50X and 400X magnification.

### Statistical analysis

Values are presented as mean ± S.D. P denotes the *p*-value. Significance was accepted for p˂0.05. Student’s paired *t* test (two tailed) and Mann-Whitney U test for comparison between two groups were carried out with GraphPad Prism 10 (GraphPad Software, Inc). Data comparing multiple experimental groups were analyzed using R4.4.3 Software (R Foundation). Data were transformed (Box-Cox) and outliers were identified by analyzing the Spearman residuals plot (absolute values >2). Normality test Shapiro-Wilk for normality, and Bartlett’s test for homocedasticity of variance. To evaluate which factors can explain the dependent variable, linear mixed-effects models controlling for the gel/kit/subject effect (random effects) were employed. Subsequently, to determine which fixed effects can explain the OD values, two complementary approaches were used based on: (i) Akaike’s Information Criterion (AIC) and (ii) hypothesis-testing by conducting an ANOVA of the model for the fixed factors. Specifically, regarding interaction, this has to be in the model in case of significant interaction and/or the interaction has to be in the model according to AIC. In the affirmative case, Tukey *post hoc* test (pairwise comparison) was performed. Both sexes were included in the analysis.

## RESULTS

### Inflamed human VECs rewire metabolism to a more energetic phenotype and exhibit upregulation of glycolytic enzymes

Our recent studies revealed that proinflammatory signals like IFN-γ and bacterial LPS rewire the metabolism of human VICs toward a glycolytic phenotype crucial for *in vitro* calcification that recapitulates the pattern observed in diseased valves (17). Given that the metabolic program of VECs has not been characterized, we performed real-time metabolic analysis of VECs derived from non-diseased valves. Under basal conditions, VECs showed higher extracellular acidification rate (ECAR), indicative of glycolysis, than VICs (Figure 1A). Exposure of VECs to IFN-γ and LPS increased respiration rate (OCR) together with ECAR reduction (Figure 1B), thus disclosing a pattern distinct from the hyperglycolysis found in inflamed VICs (17).

**Figure 1.**
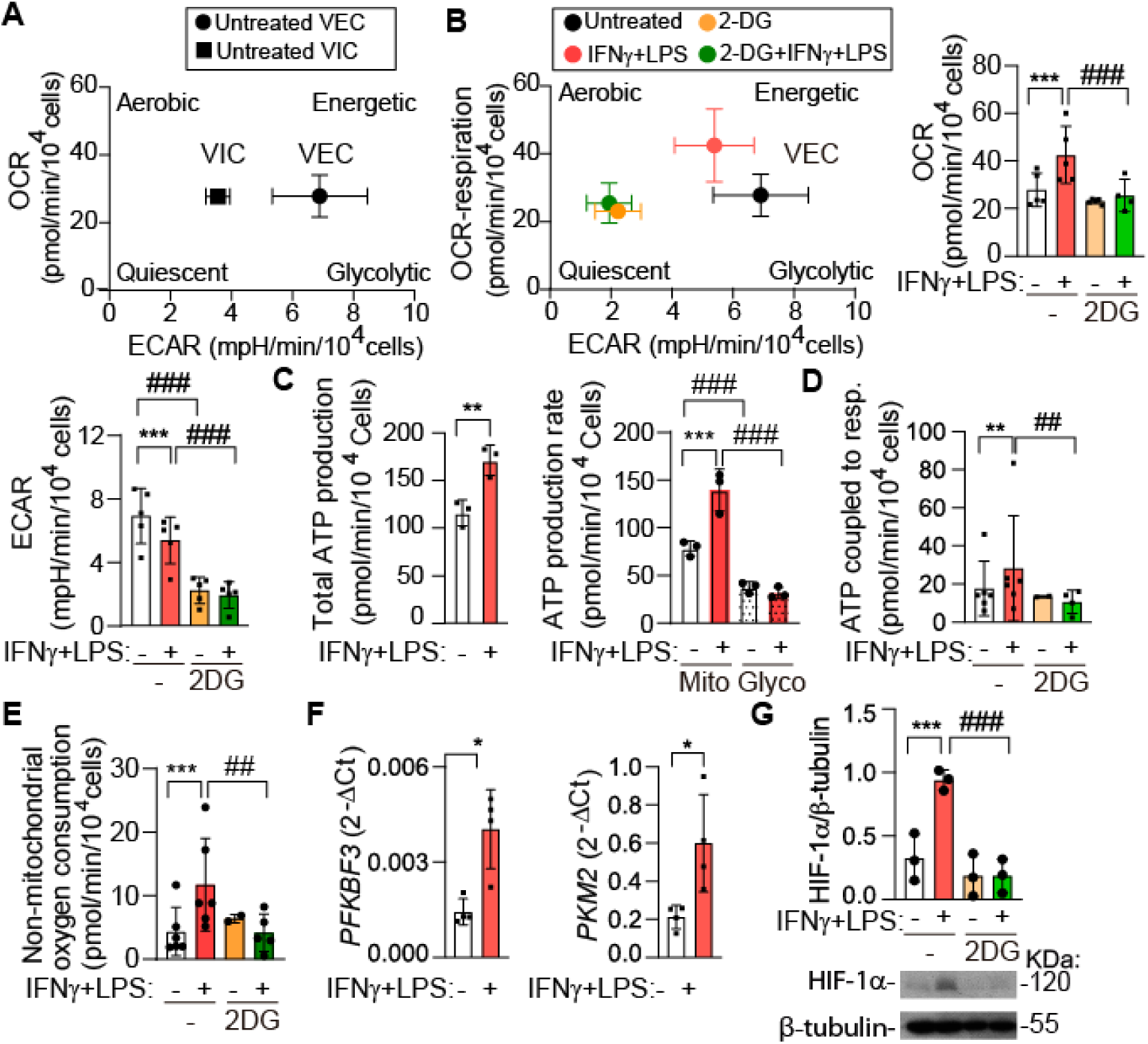
Inflammatory stimuli activate energetic metabolism and a glycolytic profile in control VECs. (**A**) Real-time metabolism of control VECs (circle) and VICs (square) under basal conditions. Energy map shows OCR vs ECAR plot, N=5 VICs and VECs isolates from independent donors. (**B-E**) Control VECs pre-incubated with 10 mM 2-DG or vehicle were activated as indicated for 24 h. Energy map, OCR, and ECAR, n=5 (**B**), Total and ATP production rate, N=3-5 (**C**), ATP-OXPHOS, N=6. (**D**), Non-mitochondrial respiration (**E**). (**F**) Gene expression by qPCR of VECs. (**G**) Western blot analysis. Representative blot and quantitation of HIF-1α protein. 2DG indicates 10 mM 2-DG. IFNγ indicates 1 µg/mL IFN-γ; LPS, 100 ng/mL LPS. Data expressed as mean ± S.D. *p<0.05, **p<0.01, ***p<0.001 compared to the control. ^##^p<0.01 ^###^p<0.001 compared to the same treatment in the other group. **B, D-E, G** Linear-mixed models (AIC) and Tukey *post hoc* (pairwise comparison). **B,** OCR, Interaction p=0.06, Activator: p<0.01, 2-DG: p<0.001; ECAR, Interaction p=0.148. Activator: p<0.01, 2-DG: p<0.001. **D**, Interaction: p=0.007, Activator: p<0.001, 2-DG p<0.001. **E,** Interaction: p<0.001, Activator: p<0.001, 2-DG p<0.01. **G**, Interaction: p<0.001, Activator: p<0.001, 2-DG p<0.001. **C** Total ATP, paired t test. ATP production rate, fixed effects from linear-mixed models. Interaction: p=0.002, Activator: p=0.24, 2-DG: p<0.001. **F**, paired t test.

ATP rate assays revealed a higher basal reliance of VECs on mitochondrial than glycolytic ATP for energy, which was further increased upon activation (Figure 1C-D). Conversely, glycolytic ATP was not altered by inflammatory agents (Figure 1C). In parallel, inflamed VECs showed increased ATP levels coupled to respiration, as well as non-mitochondrial oxygen consumption (Figure 1D-E). These effects were abrogated by pretreatment with the glycolysis inhibitor 2-deoxy-D-glucose (2-DG) (Figure 1D-E), suggesting the dependence of the process on glucose flux. Likewise, inflammatory stimuli upregulated the expression of glycolytic genes, i.e., *PFKBP3* and *PKM2* enzymes (Figure 1F), as also found in human VICs and 3D porcine VIC-VEC co-cultures (17). Further, treatment with IFN-γ and LPS stabilized the protein encoded by the glycolytic regulatory gene HIF-1α (Figure 1G). These data suggest that VECs are more glycolytic than VICs under basal conditions but exhibit a distinct shift to an energetic metabolism in inflammatory milieus, while both cell types shared a gene profile consistent with enhanced glycolysis.

### Glycolysis is required for the activation of the JAK-STAT1 signaling pathway and HIF-1α and NF-κB transcription factors and the ensuing secretion of cytokines and *PTGS2/* COX2 gene upregulation in inflamed valve cells

VICs and VECs sense inflammatory signals via the JAK-STAT system and the NF-κB route to induce the expression of glycoproteins relevant to valve pathology, like IL-6, COX-2, and adhesion molecules (6–8, 26). To address the role of glucose metabolism on the proper protein glycosylation, we used 2-DG, a non-metabolizable glucose analog and competitive inhibitor of hexokinase II, the entry-point enzyme in glycolysis (27). Western blot analysis of VICs showed that pretreatment with 2-DG blunted the stabilization of HIF-1α and the phosphorylation of STAT1 and NF-κB-p65 detected in VICs exposed to IFN-γ and LPS (Figure 2A). These findings indicate that glycolysis, at least in part, is necessary for the activation of these inflammatory routes.

**Figure 2.**
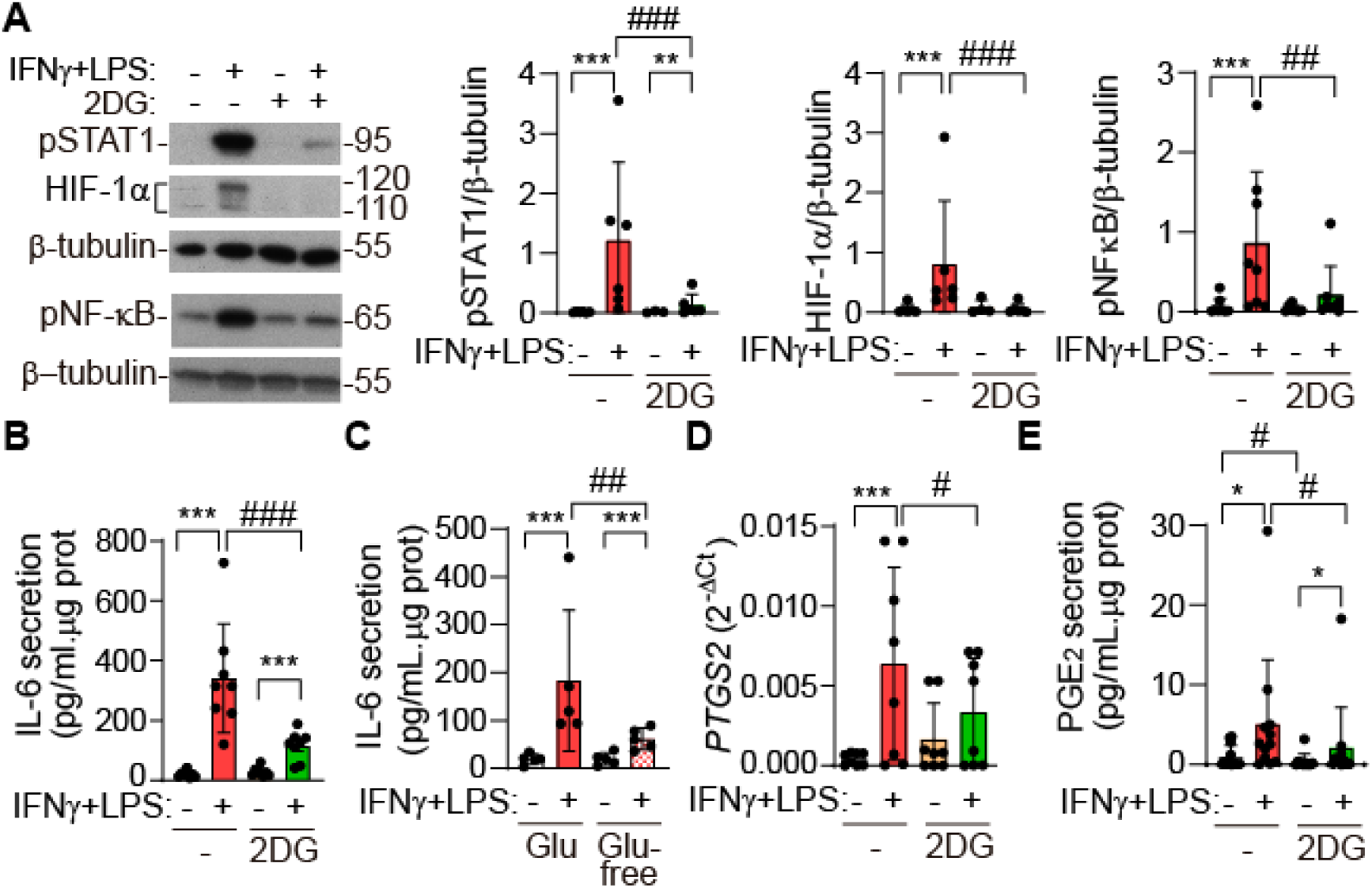
The blockade of glucose metabolism with 2-DG reduces the activation of the JAK/STAT/HIF-1α axis and NF-κB route and the subsequent secretion of IL-6 and PGE_2_ by inflamed VICs. Human VICs were pre-incubated with 10 mM 2-DG or vehicle (-) and activated for 24 h, as indicated as in Figure 1. (**A**) Western blot analysis. Representative blot and quantitation of p-STAT1, HIF-1α, and phospho-NF-κB-p65 proteins, N=6-8. (**B**, **C**, **E**). Supernatants were analyzed by ELISA for the IL-6 and PGE_2_. N=8-12. (**D**) PCR analysis of *PTSG2* (COX2) gene. Data presented as mean ± SD. *p<0.05, ***p<0.001 compared to the control. ^#^p<0.05, ^##^p<0.01, ^###^p<0.001 compared to the same treatment in the other group. **A-E** Linear mixed models; Tukey *post hoc* (pairwise comparison). **A**, pSTAT1, Interaction: p<0.01, Activator: p<0.001, 2-DG: p<0.001; HIF-1α, Interaction: p<0.001, Activator: p<0.001, 2-DG: p<0.001; p-NF-κB, Interaction: p=0.011, Activator: p<0.001, 2DG p=0.012. **B**, Interaction: p=0.002, Activator: p<0.001, 2-DG: p<0.001. **C**, Interaction p=0.013, Activator: p<0.001, Glucose: p=0.015. **D**, Interaction: p=0.015, Activator: p<0.001, 2DG p=0.39. **E**, Interaction: p=0.87, Activator: p<0.001, 2-DG: p<0.001.

Then, we investigated NF-κB-regulated inflammatory molecules relevant to CAVD pathogenesis, i.e., IL-6 and COX-2 (3). ELISA analysis revealed that 2-DG inhibited the secretion of IL-6 by inflamed VICs (Figure 2B). Likewise, glucose deprivation impaired IL-6 secretion (Figure 2C). Additionally, 2-DG prevented the upregulation of *PTGS2*/COX2 gene as well as the secretion of PGE_2_ (Figure 2D-E).

In VECs, 2-DG inhibited the stabilization of HIF-1α (Figure 1G) and the production of inflammatory mediators (Figure 3A-C). Particularly, 2-DG inhibited the secretion of IL-6 as well as the upregulation of *PTGS2*/COX2 gene (Figure 3A-B).

**Figure 3.**
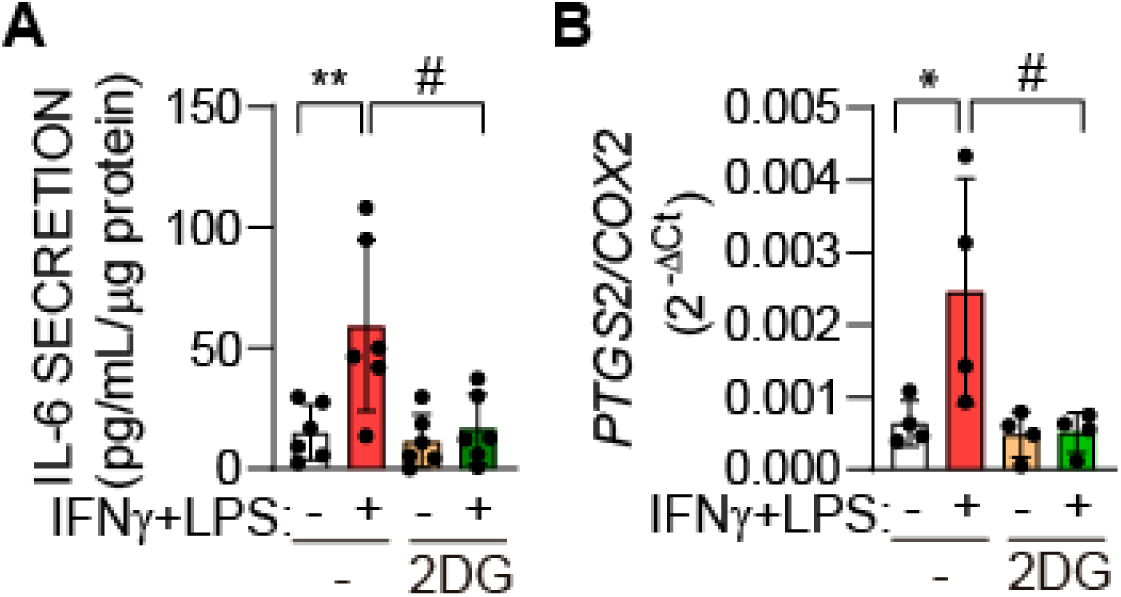
Blocking glucose metabolism with 2-DG impairs the secretion of IL-6 and *PTSG2*/COX2 gene expression in inflamed VECs. VECs were pre-incubated with 10 mM 2-DG (2DG) or vehicle (-) and activated for 24 h. (**A-B**) Cells were analyzed as in Figure 2. Data expressed as mean ± S.D. *p<0.05, **p<0.01 compared to the control. ^#^p<0.05 compared to the same treatment in the other group. N=12. **A-B**. Linear-mixed models and Tukey *post hoc* (pairwise comparison). **A**, Interaction p<0.01. Activator: p<0.001, 2-DG: p<0.001. **B**, Interaction p=0.036, Activator: p=0.92, 2-DG: p=0.64.

### Glycolysis is required for adhesion molecule maturation in inflamed VICs with ICAM-1 showing sex-dependent induction

Next, we explored the induction of VCAM-1 and ICAM-1, which are heavily glycosylated proteins regulated by NF-κB. These adhesion molecules are secreted by VICs exposed to LPS (28) and their levels are elevated in sera from patients with non-rheumatic valve disease and associated with increased prevalence and severity of aortic valve calcium in the MESA cohort (10, 11). In inflamed VICs, Western blot analysis disclosed that the induction of ICAM-1 and VCAM-1 was reduced by 2-DG (Figure 4A) and glucose deprivation (Figure 4B). Notably, 2-DG significantly decreased their induction and reduced the protein’s apparent molecular weight, as indicated by the faster migration in the gel than its full-length size counterpart (Figure 4A). This is indicative of the inhibition of their PTMs, consistent with the 2-DG disruption of N-glycosylation. Furthermore, mannose, a C-2 epimer of glucose, partly recovered 2-DG-modified ICAM-1 and VCAM-1 molecular masses and expression (Figure 4A).

**Figure 4.**
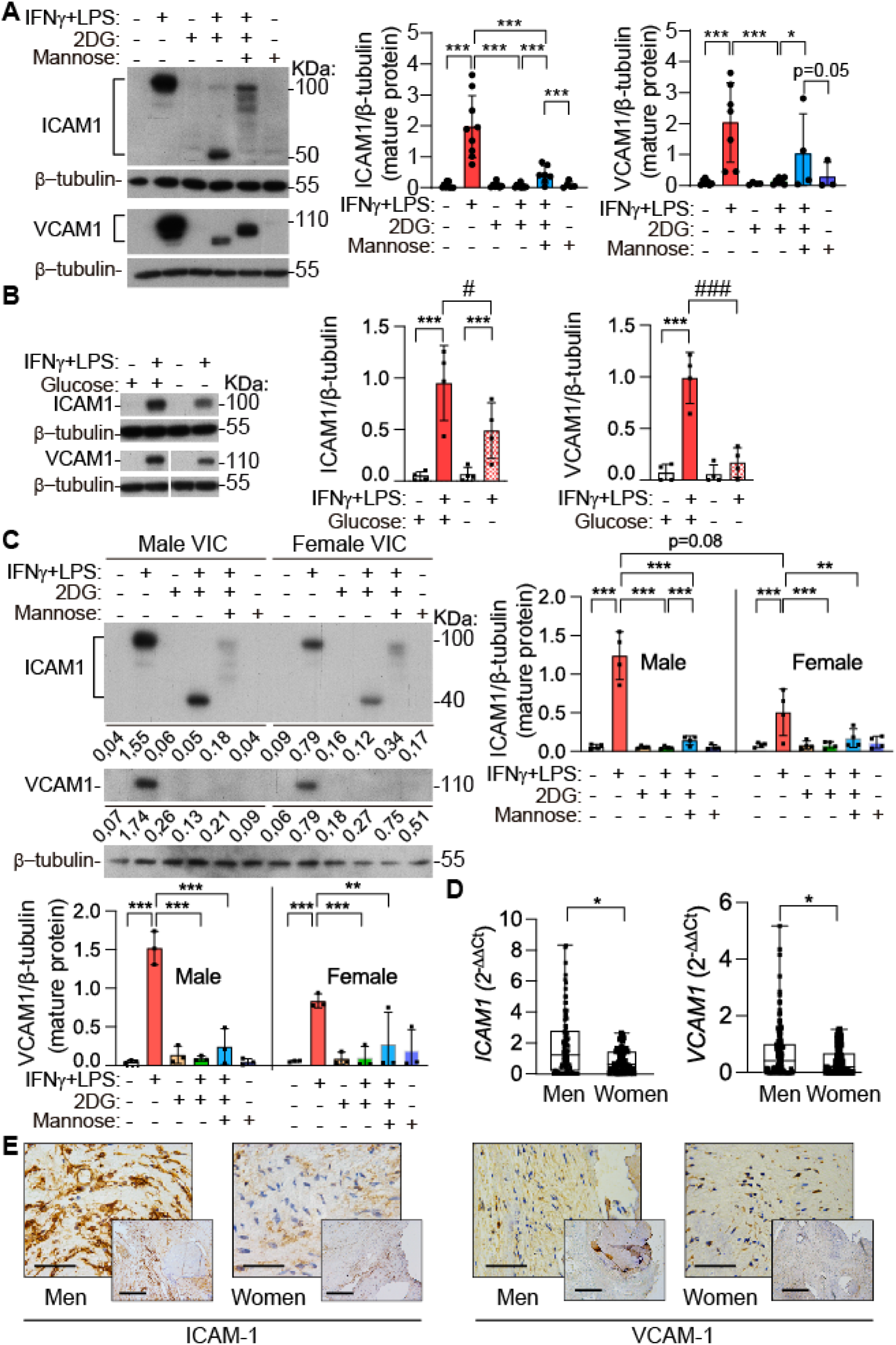
The blockade of glucose metabolism reduces ICAM-1/VCAM-1 expression and molecular mass that are partly recovered by mannose in VICs. (**A-C**) Control non-stenotic VICs were treated as indicated as in Figure 1 and ICAM-1/VCAM-1 were analyzed by Western blot. (**A**). Mannose indicates 1 mM D-(+)-mannose. The expression of the mature form, indicative of the fully processed form of the corresponding protein, was analyzed. N=7-9. (**B**) Cells were activated in media with or without glucose. N=4. (**C**) Comparison of VICs derived from male and female in the same blot. N=4 for each sex.The numbers under the gel image indicate the ICAM1/β-tubulin and VCAM1/β-tubulin ratio. (**D**) *ICAM1* and *VCAM1* mRNA expression in aortic valves from male and female patients with aortic stenosis. (**E**) Immunohistochemistry to evidence ICAM-1 and VCAM-1 expressions in aortic valves from men and women with CAVD. Scale bars, 50 and 200 µm Data expressed as mean ± S.D. *p<0.05, **p<0.01, ***p<0.001 compared to the control. ^#^p<0.05, ^##^ ^#^p<0.001 compared to the same treatment in the other group. **A** Linear-mixed one factor model factor (treatments), Tukey *post hoc* (pairwise comparison). **B** Linear-mixed models, Interaction p=0.04, Activator p<0.001, Glucose p=0.19. **C** Linear-mixed models, Tukey *post hoc* (pairwise comparison). ICAM-1 (mature form), Interaction p<0.01, Treatments p<0.001, sex p=0.62 (AIC); VCAM-1 (mature form), Interaction p<0.065, Treatments p<0.001, sex p=0.82. **D** Mann–Whitney U test, p<0.05 vs men.

Experiments comparing the effects on VICs derived from male and female donors showed similar requirement on glycolysis for adhesion molecule expression and processing in both sexes (Figure 4C). On the other hand, in every pair of male-female VICs analyzed, we observed higher induction of ICAM-1 and VCAM-1 in VICs from male as compared to female (Supplemental Tables S3-S4). Statistical analysis of Western blot data showed a tendency of higher induction of ICAM-1 in VICs derived from male than female donors (Figure 4C). The *in vivo* relevance of sex-differential induction of adhesion molecules was investigated in aortic valves explanted from CAVD patients by analyzing a cohort used in a previous study (25). PCR analysis of valve tissue revealed a higher expression of *ICAM1* gene in male as compared to female valve tissue (Figure 4D), consistent with previous data (25), and further demonstrated that *VCAM1* is expressed to a higher extent in valves from males. Moreover, aortic valves from men exhibited higher immunostaining for both ICAM-1 and VCAM-1 as compared to aortic valves from women (Figure 4E).

### Glycolysis is required for adhesion molecule expression and subsequent adhesion of leukocytes to inflamed VECs

The adhesion of monocytes to valve endothelium was then assessed given that the earliest lesion of CAVD is the presence of an inflammatory infiltrate composed of macrophages (29). Pretreatment with 2-DG in VECs inhibited the inflammation-mediated induction of VCAM-1 and ICAM-1 and reduced their molecular mass (Figure 5A), as also shown in VICs (Figure 4A). Notably, mannose did not significantly recovered 2-DG-modified glycosylated forms of ICAM-1 and VCAM-1 in VECs (Figure 5A), in contrast to VICs (Figure 4A). In addition, the upregulation of *CD62E*, which encodes the adhesion molecule E-selectin, *CD62E,* was also inhibited by 2-DG (Figure 5B).

**Figure 5.**
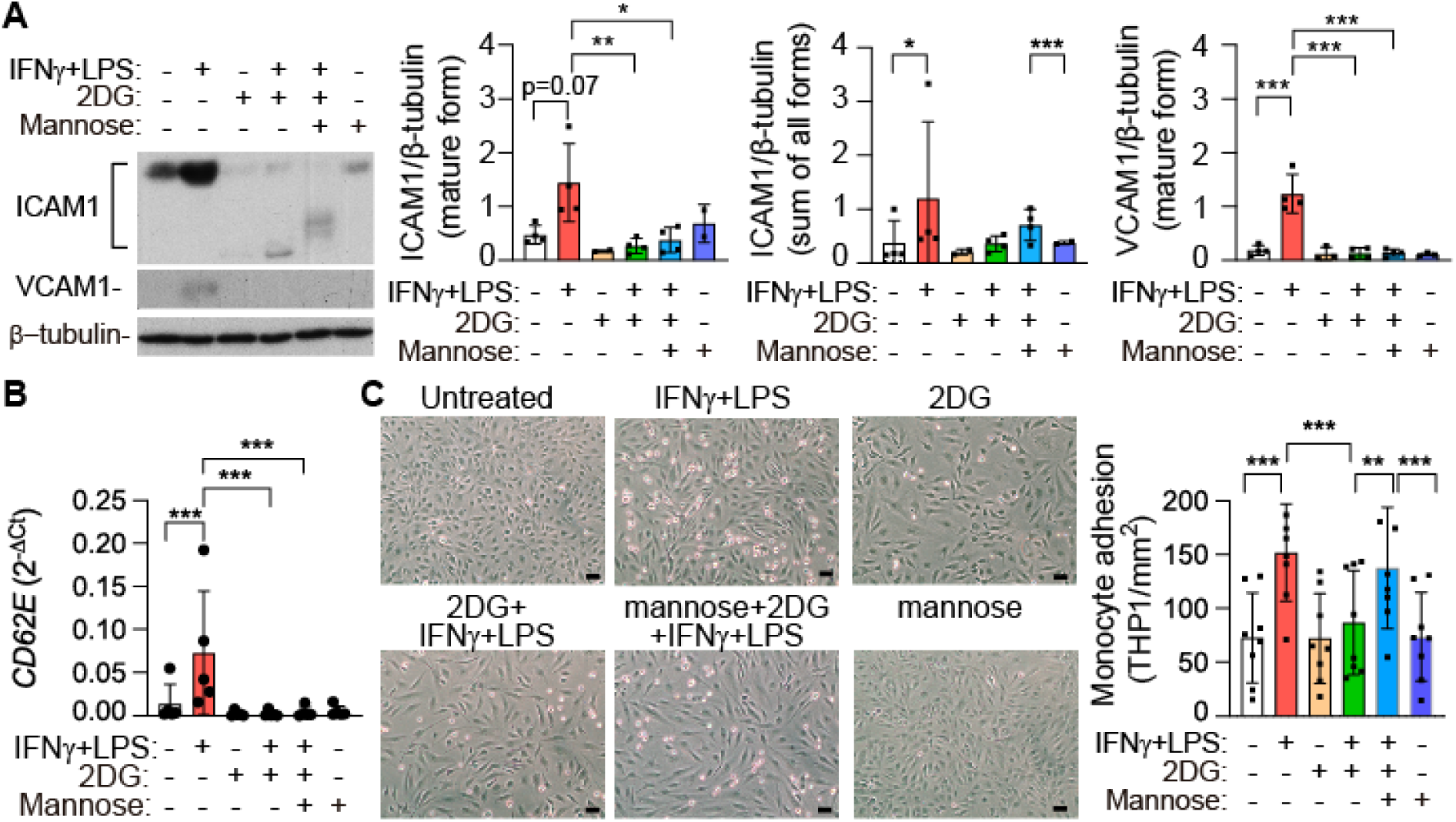
2-DG reduces the expression of adhesion molecules and the subsequent adhesion of leukocytes to inflamed VECs. Control VECs pre-incubated with 10 mM 2-DG (2DG), or vehicle were activated as in Figure 4. (**A**) VECs were collected for Western blot analysis of ICAM-1 and VCAM-1. N=4. The expression of the mature form, represented by the fully processed form of the protein, and the sum of all detected forms were analyzed. (**B**) PCR analysis of *CD62E/E-selectin* gene levels. (**C**) Adhesion assay of THP1 monocytes to VECs activated as indicated. Scale bar, 50 µm. Data expressed as mean ± S.D. *p<0.05, **p<0.01, ***p<0.001 compared to the control. **A-C** Linear-mixed one factor model (treatments), Tukey *post hoc* (pairwise comparison).

To characterize the functional implications on adhesion processes, VECs-monocyte adhesion was assayed. Quantitation of adhered monocytes showed that the number of THP1 monocytes adhered to VECs monolayers upon activation with IFNγ+LPS was markedly blunted by 2-DG, which also induced changes in VECs morphology suggestive of marked differentiation (Figure 5C). Together, data suggest that glycolysis is necessary, at least in part, for adhesion molecule production in inflamed VECs and consequent leukocyte adhesion.

### The HBP route is involved in the N-glycosylation of adhesion molecules and secretion of IL-6 in inflamed valve cells

To investigate the molecular mechanism underlying the disturbance of THP-1 monocyte adhesion elicited by 2-DG, the involvement of the HBP route was scrutinized (Figure 6A). First, we analyzed the expression of the rate-limiting enzyme glutamine fructose-6-phosphate amidotransferase (GFAT, also termed GFPT) (23). Under resting conditions, VICs express both isoforms *GFAT1*/*2*. Inflamed VICs showed upregulation of *GFAT2* levels and downregulation of *GFAT1*, *GNPNAT*, and *PGM3* transcript levels (Figure 6B). Consistent with mRNA findings, GFAT2 protein expression was increased in inflamed VICs (Figure 6C). To assess the role of the HBP route on cytokine secretion and adhesion molecule expression, we used diazo-5-oxo-L-norleucine (DON), a glutamine antagonist that inhibits GFAT. DON pretreatment reduced the secretion of IL-6 by inflamed VICs (Figure 6D). Also, DON impaired the expression of the mature form and the processing of ICAM-1 and VCAM-1 proteins in a dose dependent manner, as judged from the detection of non-fully processed forms exhibiting faster migration in SDS-PAGE (Figure 6E, Supplemental Figure S2A). Conversely, DON did not alter the expression of the sum of all detected forms (Figure 6E), suggesting the impairment of the maturation process but not protein expression. Additionally, DON did not affect GFAT2 protein expression (Supplemental Figure S2B). Data suggest that blockade of HBP prevents the incorporation of glycan moieties during glycosylation reactions as well as the maturation of adhesion molecules in VICs.

**Figure 6.**
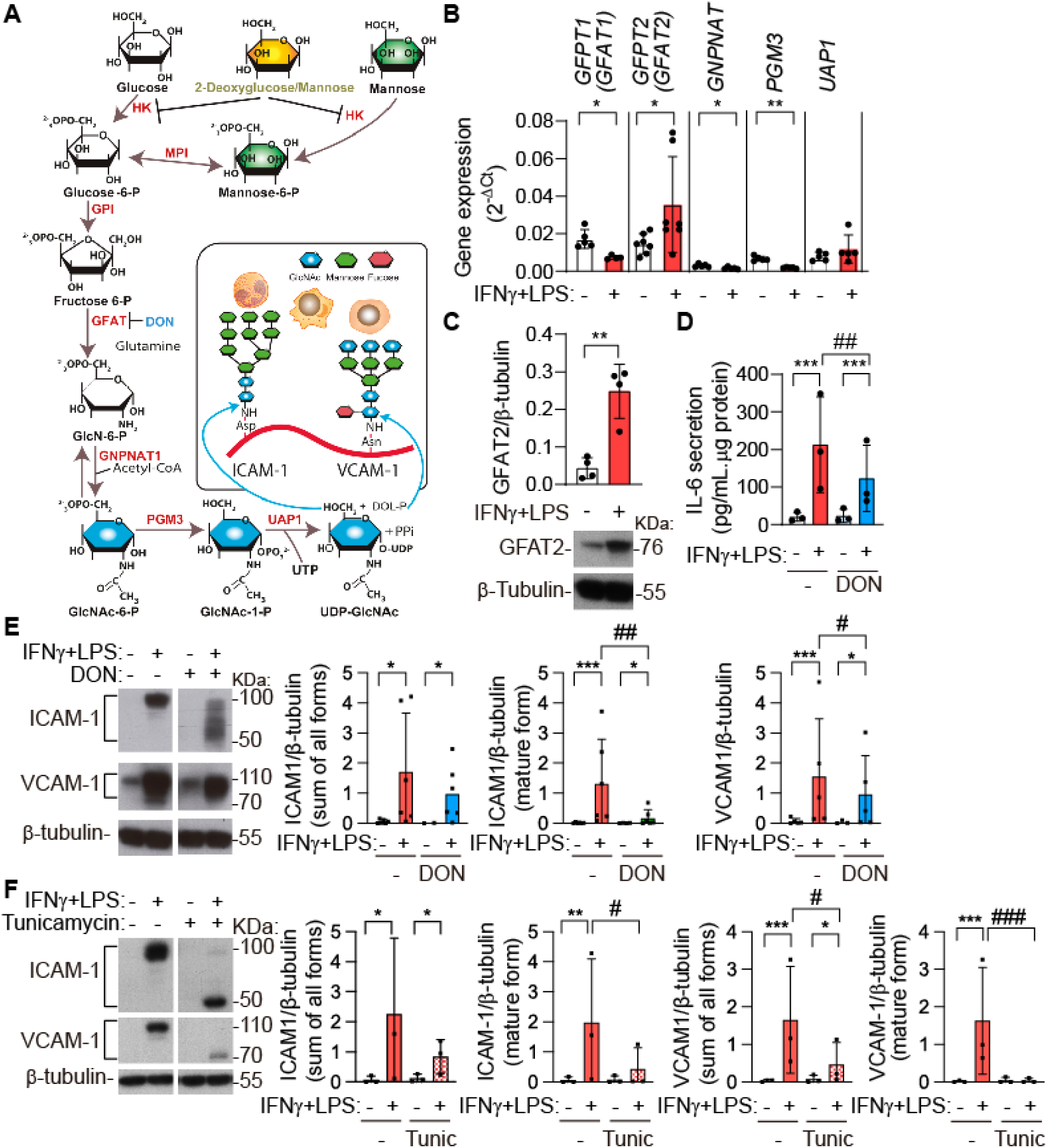
The HBP inhibitor DON reduces the N-glycosylation of adhesion molecules and IL-6 secretion in inflamed VICs. (**A**) Scheme depicting the HBP route steps and enzymes. (**B**) qPCR analysis of *GFPT1/2, GNPNAT, PGM3 and UAP1* genes. (**C**) Western blot analysis of the GFAT2 protein. N=4. (**D-F**) VICs were pretreated as indicated and activated for 24 h and then analyzed by ELISA, N=6 (**D**) or Western blot, N=3-5 (**E-F**). DON, 100 μM DON, Tunicamycin, 100 ng/mL tunicamycin. Data expressed as mean ± S.D. *p<0.05, **p<0.01, ***p<0.001 compared to the control. #p<0.05 compared to the same treatment in the other group **B-C** Paired t test. **D-F** Linear-mixed models, Tukey *post hoc* (pairwise comparison) **D**, Interaction: p<0.001, Activator: p<0.001, DON: p<0.001; **E** ICAM-1 all forms, Interaction: p=0.46, Activator: p<0.001, DON: p=0.96; ICAM-1 mature form, Interaction: p<0.01, Activator: p<0.001, DON: p<0.01; VCAM-1, Interaction: p=0.03, Activator: p<0.001, DON: p=0.12. **F** ICAM-1 all forms, Interaction: p=0.44, Activator: p<0.001, Tunicamycin: p=0.76; ICAM-1 mature form, Interaction: p=0.03, Activator: p<0.001, Tunicamycin: p=0.72; VCAM-1 all forms, Interaction: p<0.01, Activator: p<0.001, Tunicamycin: p=0.064 (AIC); VCAM-1 mature form, Interaction: p<0.001, Activator: p<0.001, Tunicamycin: p<0.001.

Tunicamycin, an inhibitor of UDP-GlcNAc:dolichylphosphate GlcNAc-1-phosphotransferase that catalyzes the first step in the biosynthesis of N-glycans utilizing UDP-GlcNAc as substrate (30), led to drastic reduction in both fully glycosylated ICAM-1 and VCAM-1 proteins (Figure 6F) mimicking the effect of 2-DG (Figure 3A) and consistent with the impairment of N-glycosylation.

DON reduced the fully glycosylated form of ICAM-1 in a dose-dependent manner, while did not alter the sum of all isoforms (Figure 7B, Supplemental Figure S1C), suggesting the inhibition of the maturation but not the expression of ICAM-1.

**Figure 7.**
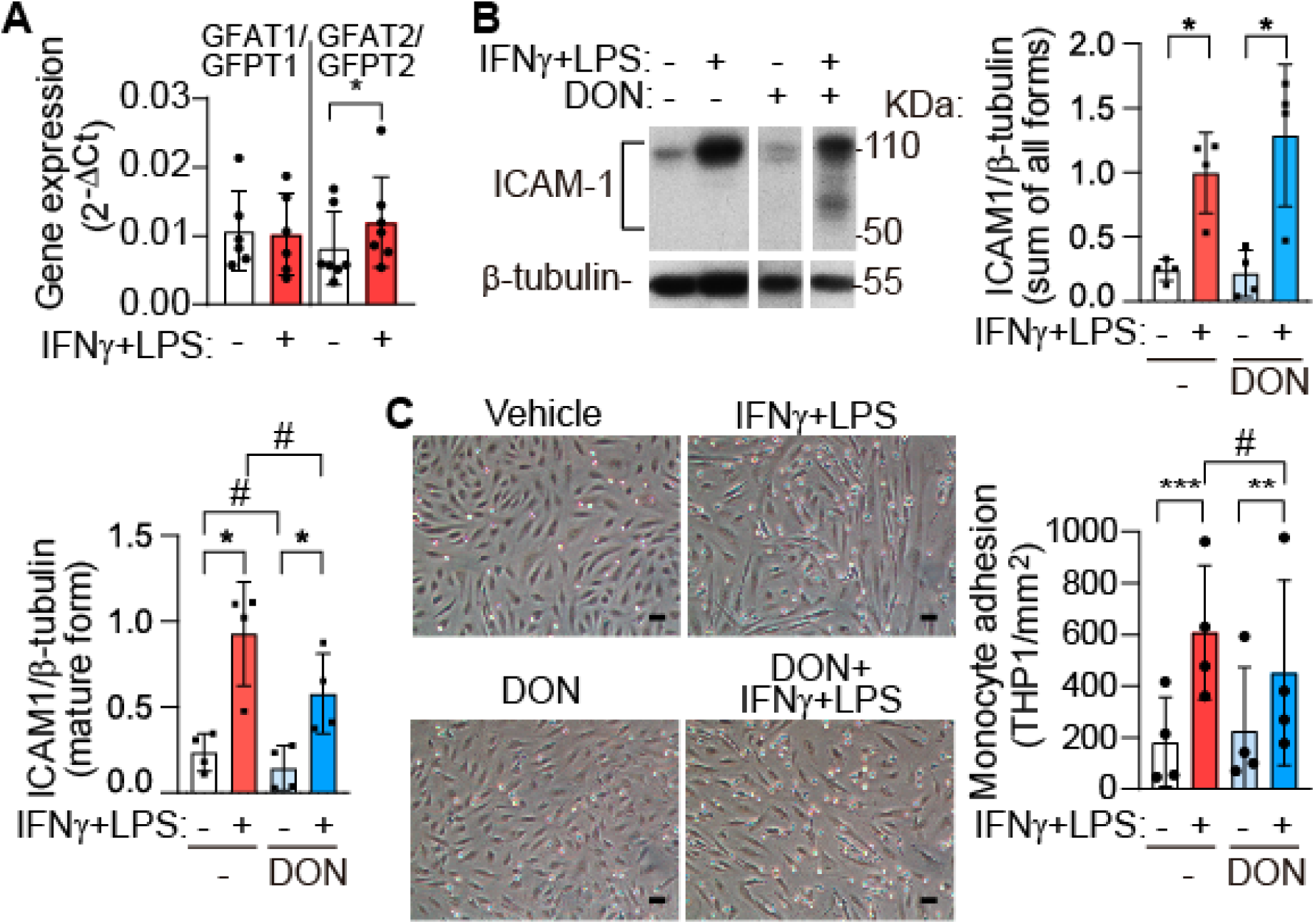
The HBP inhibitor DON reduces the glycosylation of ICAM-1 and the adhesion of monocytes in inflamed VECs. Human VECs were pretreated and activated for 24 h, as indicated. (**A**) qPCR analysis of *GFPT1/2* genes. (**B**) Western blot analysis of ICAM-1. (**C**) Adhesion of THP1 monocytic cells to VECs monolayers. Scale bar, 50 µm. DON, 100 μM DON. Data expressed as mean ± S.D. *p<0.05, **p<0.01, ***p<0.001 compared to the control. #p<0.05 compared to the same treatment in the other group. **A**, paired t test. **B-C** Linear mixed models, **B** The sum of all form of ICAM-1, Interaction p=0.48, Activator: p<0.001, DON: p<0.001; Interaction p=0.21, Activator: p<0.001, DON: p=0.76, mature form of ICAM-1. **C** Monocyte adhesion, Interaction p=0.02, Activator: p<0.001, DON: p=0.24 Tukey *post hoc* (pairwise).

Alike VICs, quiescent VECs express isoforms *GFAT1* and *GFAT2*, while inflamed VECs showed upregulation of *GFAT2* mRNA expression (Figure 7A). Also, DON reduced the fully glycosylated form of ICAM-1 in a dose-dependent manner, while did not alter the sum of all isoforms (Figure 7B, Supplemental Figure S1C). Furthermore, DON partly inhibited the adhesion of monocytes to VECs monolayers (Figure 7C). These results suggest that blockade of HBP prevents the glycosylation and maturation of ICAM-1 and the subsequent monocyte-VECs adhesion. Together, these data indicate the involvement of HBP in the glycosylation of adhesion molecules in aortic valve cells exposed to inflammatory agents and suggest a role of GFAT2 in view of its induction by inflammatory stimuli.

## Discussion

Our study discloses the role of glycolysis and its offshoot pathway HBP in the production of mature glycoproteins implicated in CAVD pathogenesis. Our findings unveil that disrupting sugar-driven PTMs leads to adverse consequences in aortic valve cells, because they play a key role in the proper function of leukocyte adhesion molecules and other proteins involved in the inflammatory response. Therefore, these PTMs are essential for the recruitment of cells of the monocytic lineage to aortic valve leaflets in the initial stages of CAVD (27).

An unexpected finding has been the distinct energetic metabolism of the different types of resident valve cells in both basal and inflammatory conditions. Basally, VECs are more glycolytic than VICs, and rely more on mitochondrial than glycolytic ATP for energy. Inflamed VECs exhibit an energetic metabolism based on mitochondrial ATP production with no change in glycolytic ATP. By contrast, in a recent report on human VICs we showed a metabolic rewiring toward glycolysis in inflammatory settings that recapitulates the metabolic pattern observed in diseased valves (17). Notably, both valve cells share a glycolytic gene signature in inflammatory environments, wherein glycolysis may be relevant for detrimental consequences. Endothelial cells use O_2_ for other crucial functions besides ATP production, for instance, NO production by cytosolic endothelial NO synthase. Since aortic valve endothelium is exposed to the combined action of hyperoxic 21% O_2_ and shear-stress, a strong OXPHOS activity may be necessary to counter the untoward effects associated with shear-induced superoxide production and the ensuing generation of peroxynitrite, which ultimately may suppress respiration by irreversibly inhibiting NADH-ubiquinone reductase (31). Moreover, a recent study has shown that a mitochondrial ATP supply is necessary for calcium-dependent, nitric oxide-mediated endothelial control of vascular tone (32). On the other hand, VECs exposed to inflammatory stimuli may display a pro-glycolytic profile as part of a broader metabolic adaptation to ensure a steady supply of intermediates required for branching pathways supporting antioxidant defense, protein glycosylation, and nucleotide biosynthesis (33). Thus, a pro-glycolytic profile may occur even in the absence of measurable extracellular acidification, reflecting a metabolic shift oriented toward sustaining anabolic and signaling demands rather than lactate accumulation.

Our study also discloses in VECs specific differences with endothelial cells from other organs. Quiescent VECs show commonalities with endothelial cells from large vessels and differences with microvascular endothelial cells as regards fuel dependence (34). The metabolic rewiring of endothelial cells in response to external signals seems to differ according to their origin. VECs dependence on oxidative phosphorylation increases under inflammatory conditions. However, since the available studies focused on vascular cells activated with FGF, VEGF, or mechanical stress and showed increased glycolytic fluxes, differences might be attributed to the type of stimuli rather than tissue specificity.

Disrupting glucose metabolism in inflamed VICs significantly impacts JAK-STAT-, HIF-1α- and NF-κB-dependent gene expression. NF-κB, a master regulator of inflammation, has been identified as an essential molecular regulator for both VECs and VICs participation in CAVD pathogenesis (35). Our data showed HIF-1α stabilization and the subsequent upregulation of glycolytic enzymes in VECs in inflammatory settings. Mechanistic cell-specific differences have been found in endothelial cells exposed to IFN-γ. In coronary artery endothelial cells, glycolysis inhibition by IFN-γ was mediated by HIF-1α destabilization (36), while in valvular endothelial cells co-stimulation with IFN-γ and LPS promoted HIF-1α stabilization and subsequent transcriptional upregulation of glycolytic enzymes.

Our findings unveil an association between glycolysis and inflammation on the expression and maturation of glycoproteins relevant to valve disease. IFN-γ and pathogen molecular patterns generate protein and lipid inflammatory mediators that propagate inflammation. Glycolysis is crucial for the PTMs necessary for glycoprotein maturation in resident valve cells. Pharmacological blockade with 2-DG and glucose-deprivation studies support the role of glucose metabolism beyond its role in energy production. It should be noted that 2-DG inhibits the entry-point enzyme in glycolysis (27), therefore its contribution of other offshoot routes of glucose metabolism cannot be ruled out. It could also be argued that 2-DG may exert nonspecific effects associated with endoplasmic reticulum stress and the ensuing unfolded protein response (37); however, these effects also depend on hexosamine metabolism.

In inflammatory settings, glycolysis is required for the secretion of IL-6, a cytokine associated to CAVD in both VIC and VEC populations. This cytokine is highly expressed in severe CAVD and is a strong inducer of *in vitro* calcification (3, 38). Our findings also highlight the role of glycosylation in COX-2 expression in resident valve cells. Glycolysis seems to be required for the induction and proper activity of COX-2 in VICs exposed to an inflammatory milieu. Our findings are consistent with previous evidence in rabbit articular chondrocytes, where 2-DG reduces COX-2 expression and N-glycosylation as well as COX-2 activity via a Src kinase-dependent pathway (39). Notably, high expression of COX-2 has been found in calcified areas of aortic valve tissue (9) and a recent study identified several eicosanoids in plasma and valve tissue from CAVD patients. Moreover, COX–derived prostanoids showed a marked correlation with disease severity (40). Of particular relevance to CAVD are PGE_2_ and PGF_2α_ due to their involvement in the process of osteogenesis, which plays a crucial role in the development of the disease (40).

Glycosylations dependent on the HBP are critical for the maturation of many proteins (41). Enzyme-catalyzed chemical modifications of proteins with sugars involves from monosaccharides to complex structures that are classified as N-, O-, or C-linked glycosylations, depending on the amino acid residue they are attached to. Adhesion molecules relevant to CAVD like ICAM-1 and VCAM-1 are heavily glycosylated proteins, including N-acetyl-D-glucosamine and mannose, and also fucose in the case of VCAM-1 (42, 43), and play a role in the process of leukocyte adhesion and transendothelial migration (44). Our findings show that inhibition of N-linked glycosylation inhibitor by tunicamycin and 2-DG reduce the inflammation-triggered expression and the molecular mass of ICAM-1 in inflamed VICs, suggesting that ICAM-1 produced by inflamed VICs include the N-glycosylated mature form. VCAM-1 protein shift is less apparent than the observed with ICAM-1. Notably, mannose partly restores the mature glycosylated form of adhesion molecules in inflamed VICs but not in VECs, arguing for valve cell-dependent effects. To note, the pattern of glycosylation restoration of ICAM-1 by mannose differs in both types of valve cells, with glycoforms showing higher molecular weight in inflamed VICs than in VECs. This agrees with the notion that ICAM-1 glycosylation depend on cell type and activation state, and the well-known distinct functions of different N-glycoforms (45, 46). Based on this, one might speculate that VICs and VECs may have variations in ICAM-1 glycosylation that might affect its function and interaction with other molecules.

Notably, HBP plays a dual role in cardiac pathophysiology and has emerged as a novel therapeutic target of cardio-metabolic diseases (47). While acute activation confers cardioprotection, chronic activation promotes the initiation and progression of diseases. Consistent with this notion, GFAT2 is overexpressed in valve cells under exposure to inflammatory agents previously associated with valve cell calcification. Blockade with DON prevented the glycosylation of ICAM-1 in a dose dependent manner. GFAT activity has recently been associated with cardiovascular diseases since GFAT2, the major heart isoform, is increased in response to hypertrophic stimuli and mediates cardiac hypertrophy through HBP-O-GlcNAcylation-Akt pathway (48). Candidates for the regulation of GFAT2 in inflamed valve cells may include HIF-1α and XBP-1 (reviewed in (47)).

An interesting finding is the sex-differential induction of ICAM-1 elicited by inflammatory stimuli, with VICs derived from men showing higher levels than VICs from women. These differences are relevant to valve disease pathogenesis since adhesion molecules are expressed to a higher extent in stenotic valve tissue. Results are consistent with previous data demonstrating higher levels of ICAM-1 in diseased valves explanted from men (25) and further demonstrate that VCAM-1 also showed sex-differential expression in diseased valves. On the other hand, data show similar processes of protein maturation in VICs derived from males as compared to females.

### Limitations of the study

The number of valves from women available for this study was limited, although it is important to emphasize that adhesion molecule data were consistent, and sex differences on the induction of ICAM-1 and VCAM-1 were detected in every pair of male-female VICs chosen randomly. Moreover, sex differences were also validated in *ex vivo* studies performed in aortic valves from CAVD patients obtained after aortic valve replacement. Also, the limitations of 2D cellular models stand from the culture on plastic plates that enriches the myofibroblast phenotype in the case of VICs.

### Conclusions

Glycolysis and the side-branch route hexosamine biosynthesis pathway are necessary for nutrient-driven post-translational modifications of mediators in inflamed valve cells and the subsequent leukocyte adhesion, a process relevant to initial stages of CAVD.

## Data availability

Datasets have been deposited to the Digital.CSIC repository (institutional repository of the Spanish National Research Council. Data will be made available upon reasonable request https://doi.org/10.20350/digitalCSIC/17797

## Supplemental Material

https://doi.org/10.20350/digitalCSIC/17800

Supplemental Figures S1–S2

Supplemental Tables S1–S4

Major resources table

Original Uncropped Gels

## Acknowledgements

The authors are grateful to the patients and medical personnel from the Hospital Clínico de Valladolid for their invaluable cooperation. Graphical abstract was created with BioRender.com.

## AUTHOR CONTRIBUTIONS

Conceived and designed research, T.S.B., M.S.C, M.P.R, and C.G.R.

Performed experiments, T.S.B, M.P.R., and C.G.

Analyzed data, T.S.B., M.P.R, E.P-R, N.L.A, J.L., A.S.R, and C.G.R.

Interpreted results of experiments T.S.B., M.P.R, M.S.C., N.F., N.L-A, and C.G.R.

Prepared figures T.S.B., M.P.R, N.L-A, and C.G.R..

Drafted manuscript, C.G.R.

Edited and revised manuscript, M.S.C, T.S.B., M.P.R, N.F., J.L., A.S.R, E.P-R, and N.L.A.

Approved final version, M.S.C, T.S.B., M.P.R, N.F., J.L., A.S.R, E.P-R, C.G, and N.L.A.

## Non-standard Abbreviations and Acronyms

AIC: Akaike’s Information Criterion
CAVD: Calcific aortic valve disease
COX-2: Cyclooxygenase 2
GFAT: Glutamine fructose-6-phosphate amidotransferase
HBP: Hexosamine biosynthesis pathway
HIF-1 α: Hypoxia inducible factor 1α
IFN-γ: Interferon-γ
IL-6: Interleukin 6
LPS: Lipopolysaccharide
JAK: Janus kinase
ICAM-1: Intercellular adhesion molecule-1
PGE_2_: Prostaglandin E_2_
PTMs: Post-translational modifications
STAT: Signal transducer and activator of transcription
UDP-GlcNAc: Uridine diphosphate N-acetylglucosamine
VCAM-1: Vascular cell adhesion molecule
VECs: Aortic valve endothelial cells
VICs: Aortic valve interstitial cells
2-DG: 2-deoxy-D-glucose

